# Phylogenetic tree inference from local gene content

**DOI:** 10.1101/017699

**Authors:** Galina Glazko, Michael Gensheimer, Arcady Mushegian

## Abstract

**Background:** Complete genome sequences provide many new characters suitable for studying phylogenetic relationships. The limitations of the single sequence-based phylogenetic reconstruction prompted the efforts to build trees based on genome-wide properties, such as the fraction of shared orthologous genes or conservation of adjoining gene pairs. Gene content-based phylogenies, however, have their own biases: most notably, differential losses and horizontal transfers of genes interfere with phylogenetic signal, each in their own way, and special measures need to be taken to eliminate these types of noise.

**Results:** We expand the repertoire of genome-wide traits available for phylogeny building, by developing a practical approach for measuring local gene conservation in two genomes. We counted the number of orthologous genes shared by chromosomal neighborhoods (“bins”), and built the phylogeny of 63 prokaryotic genomes on this basis. The tree correctly resolved all well-established clades, and also suggested the monophily of firmicutes, which tend to be split in other genome-based trees.

**Conclusions:** Our measure of local gene order conservation extracts strong phylogenetic signal. This new measure appears to be substantially resistant to the observed instances of gene loss and horizontal transfer, two evolutionary forces which can cause systematic biases in the genome-based phylogenies.

## Background

There are about 2.5×10^113^ possible topologies for the unrooted tree of 63 species (the number of prokaryotes in the NCBI COG database in early 2004), and the correct topology is not known. Sequence-based phylogenies of prokaryotes may differ from one another, depending on which sequences are chosen. The incongruent tree topologies have to be explained by a combination of differential gene losses, unrecognized gene paralogy [1], or by variation of evolutionary rates among genes and among species, which may cause long-branch attractions and related tree distortions [2]. Horizontal gene transfer (HGT), previously thought to be inconsequential, may in fact be another significant factor in evolution of protein-coding genes in prokaryotes, obscuring phylogenetic signal (discussed in references [1] and [3]). rRNA-based phylogeny is also not exempt from artifacts, as unequal evolutionary rates and HGT could have played a role in rRNA evolution as well [4].

One way to address the phylogeny problem is to continue using sequence data, but try to eliminate these artifacts. Another way is to use genomic traits other than sequence alignments. Several such traits to study microbial genome evolution have been proposed in recent years, mostly derived from the conservation of gene content in completely sequenced genomes [5, 6].

Genome content-based traits are complementary to gene sequences, in a sense that they are differently impacted by evolutionary forces, and therefore may be less prone to some of the noises from which the sequence alignments suffer. Genome-content trees, however, have their own biases. Most notably, the bacteria with parasitic lifestyle are sometimes placed together into artificial clades, even though similarities of gene sequences would have positioned these parasitic genomes in the separate clades, close to their respective free-living relatives [6, 7]. The most likely explanation for such bogus clades is convergent evolution of gene content due to parallel gene loss [6], where distinct bacteria have independently lost many of the same biosynthetic enzymes while adapting to the nutrient-rich environments of the host. Genome size normalization has emerged as an important way to correct for this bias [8], but the most appropriate way to normalize is still under debate [9]. HGT-related artifacts may affect genome-based trees as well (discussed in reference [10]) and in such cases, one has to detect “phylogenetically discordant sequences” (PDS), i.e., those sequences which display abnormal phylogenies, and remove them from consideration in order to minimize these artifacts [11].

We present a new measure of similarity between genomes, based on counting genes that belong to conserved gene bins, i.e., to the chromosome segments sharing large number of orthologous genes, not necessarily in the same order. It is known that the extent of gene synteny between genomes decreases with evolutionary distance [12], with only a small number of operons remaining conserved in all prokaryotes [13]. Conserved gene bins include more genes than conserved operons, and we show that bins contain signal strong enough as to be suitable for resolving both close and more distant phylogenetic relationships. Moreover, the intergenome distance measure based on the number of genes in bins does not seem to be particularly sensitive to gene loss and HGT.

## Results

### Comparison of gene bins in two genomes

Our approach is to consider chromosome segments (“bins”) of constant size in two genomes, and to seek a pairing between bins in two genomes, where each bin in the smaller genome is matched up with a unique bin in a larger genome. Consider genomes 1 and 2 and an ordered pair of bins. The *i*th and *j*th bins *b*_*i*1_ and *b*_*j*2_ of size *k* can share from 0 to *k* genes (we used NCBI COGs instead of genes – see Methods and Discussion for details), and we want to find, for each bin in a smaller genome, such a matching bin in the larger genome, that two bins share as many genes as possible, up to *k*, ignoring the order of genes within bins. The value that is maximized in this way, for each bin in smaller genome, is called bin score, *s*_*12*_=max_*ij*_{*b*_*i*1_∩*b*_*j*2_} where *i* and *j* run over the number of bins in two genomes. Two high-scoring bins contain a conserved (maybe rearranged with indels) group of orthologous genes/COGs. Simultaneously, we want to obtain a global bin assignment between two genomes by maximizing the sum of bin scores; this value is called global assignment score, *Gs*. Global assignment score shows how many genes in two genomes are assigned after bin matching is completed, and its maximum is equal to the size of the smaller genome.

The optimal assignment of bins from two genomes is an instance of the fundamental combinatorial problem (generalized assignment problem, GAP), best-studied in the context of such applications as job scheduling, routing and facility location. GAP is the problem of finding the minimal cost assignment of jobs to machines such that each job is assigned to exactly one machine, subject to capacity restrictions on the machines. In our case bins in smaller genome correspond to jobs, bins in larger genome - to machines, and we are looking for the assignment which maximizes the global score. Due to NP-hardness of GAPs, recent papers tend to limit the optimization tasks to the space of less than 1,000 binary variables [14]. The size of our task is at least three times larger than this boundary, so we sought a compromise between time complexity and the evolutionarily meaningful *k* (which results in thousands of bins per genome), and designed a fast heuristic algorithm, which finds a global assignment in an iterative mode.

### Pairwise genomes assignments

Using the algorithm described in Methods, we produced 1953 pairwise assignments of 63 genomes. Global assignment scores were also computed for 100 replicates of genomes with jumbled gene order in each pair; the random probability of obtaining the score same or higher than the observed one was always ≤10^-5^. For each pair of genomes *G*_*i*_, *G*_*j*_, we collected all bins of length 10 sharing more than one COG. The maximum number of bins (399) was shared by two *E.coli* strains (Figure 1A; these genomes also had the largest global score, with 3207 out of 3985 COGs found in bins). The smallest score of 27 (27 COGs in 9 bins; Figure 1B) was observed for the pair *Methanopyrus kandleri* (Mka) - *Mycoplasma pulmonis* (Mpu). In general, the number of genes in bins is about 10% larger than the number of conservative adjacent gene pairs. For example, for pairs of archaeal genomes, the average number of conserved adjacent gene pairs is 267 [6] and the average number of genes in bins is 291 (this study).

**Figure 1.**
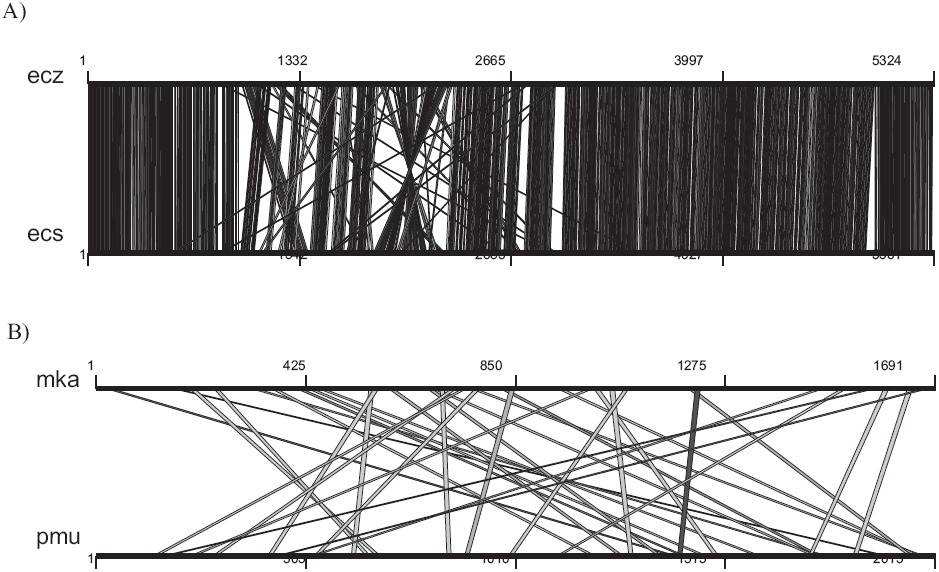
Pairwise assignment of gene bins: (A) two strains of *E.coli*. (B) archaeon *Methanopyrus kandleri* and bacterium *Mycoplasma pulmonis*.

### Distance measure

We normalized the total number of genes in bins (global assignment score, *Gs*), by the weighted average genome size, 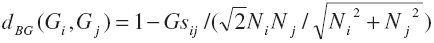. As shown recently [11], this distance is more resistant to the difference in genome sizes than the alternatives, such as division by geometric average of genome sizes or by the smallest of two genomes. A tree was inferred based on the *d*_*BG*_ distance using standard neighbor-joining (NJ) algorithm. The statistical support for internal nodes was obtained by delete-half-jackknife method [15], i.e., randomly selecting 50% of bins and recalculating the trees over 100 replications [5]. The branch support in the tree varied from 21 to 100 percent, and we discuss only internal nodes with support of 50 percent or higher.

### Phylogenetic hypothesis and comparisons with other phylogenetic trees

Phylogenetic tree of complete bacterial and archaeal genomes, obtained using *d*_*BG*_ distance measure and NJ algorithm, is shown in Figure 2. We compared this tree (Tree8, Table 1 and Figure 2) with seven other phylogenetic reconstructions:

- Two trees based on sequence similarity between aligned orthologous proteins:
  - ML (maximum likelihood) tree of 32 concatenated ribosomal proteins (Figs. 6, 7 in Wolf et al. [6]), called Tree1 in the sequel and in Table 1;
  - Fitch-Margoliash tree based on the normalized BLASTP scores (Fig. 5 in Clarke et al. [10]), Tree2;
- Three trees based on orthologous genes or gene families:
  - NJ supertree, based on a supermatrix, made by concatenation of binary matrices of orthologous gene families, where for the nodes having more than 50% bootstrap support, all the genes linked by an internal tree branch are coded as 1, the other genes being coded as 0. (Fig. 4A in Daubin et al. [16]), Tree3;
  - NJ tree based on the gene content (Fig. 1 in Korbel et al. [8]), Tree4;
  - NJ tree based on the gene content with weighted characters (Fig. 4 in Dutilh et al., [11]), Tree5;
  - maximum parsimony tree based on the presence-absence of gene families (Fig. 1 in House et al. [17]), Tree6;
- One tree based on the chromosomal proximity of orthologous genes:
  - Dollo parsimony tree based on the adjoining gene pairs (Fig. 4 in Wolf et al. [6]), Tree7.

**Table 1.** Evolutionary hypotheses suggested by the genome-wide phylogenies

| Hypothesis | Tree<sup>a</sup>1 | Tree2 | Tree3 | Tree4 | Tree5 | Tree6 | Tree7 | Tree8 |
| --- | --- | --- | --- | --- | --- | --- | --- | --- |
| 1. Monophyly of two domains, Archaea and Bacteria | + | + | + | + | + | + | + | + |
| 2. Separation of low and high GC Gram-positive bacteria | + | + | + | + | + | + | + | + |
| 3. Paraphyly of Euryarchaeota | + | + | - | + | + | + | + | + |
| 4. Chlamydia/Spirochetes clade | + | + | - | - | + | - | + | + |
| 5. Thermotoga is clustering with Gram-positive bacteria<sup>b</sup> | - | - | - | + | + | + | + | + |
| 6. Aquifex is clustering with proteobacteria<sup>c</sup> | - | - | - | + | + | - | + | + |
| 7. Monophyly of Gram-positive bacteria | - | - | - | - | - | - | - | + |
| 8. Cyanobacteria-Deinococcus-Actinomycetales clade<sup>‡</sup> | + | - | - | + | - | - | - | - |

a-Tree1: maximum likelihood tree of 32 concatenated ribosomal proteins [6]; Tree2: Fitch-Margoliash tree based on the normalized BLASTP scores [10]; Tree3: NJ supertree, based on a supermatrix [16]; Tree4: NJ tree based on the gene content [8]; Tree5: NJ tree based on the gene content with weighted characters [11]; Tree6: maximum parsimony tree based on the presence-absence of gene families [17]; Tree7: Dollo parsimony tree based on the adjoining gene pairs [6]; Tree8: this work.
b-In tree 3, there is the Thermotoga-Aquifex clade, basal to Proteobacteria.
c-The two lineages are the deepest among bacteria in Tree 8, but no specific sister relationship is evident; actinomycetes group with other Gram-positive bacteria in Tree 8.

**Figure 2.**
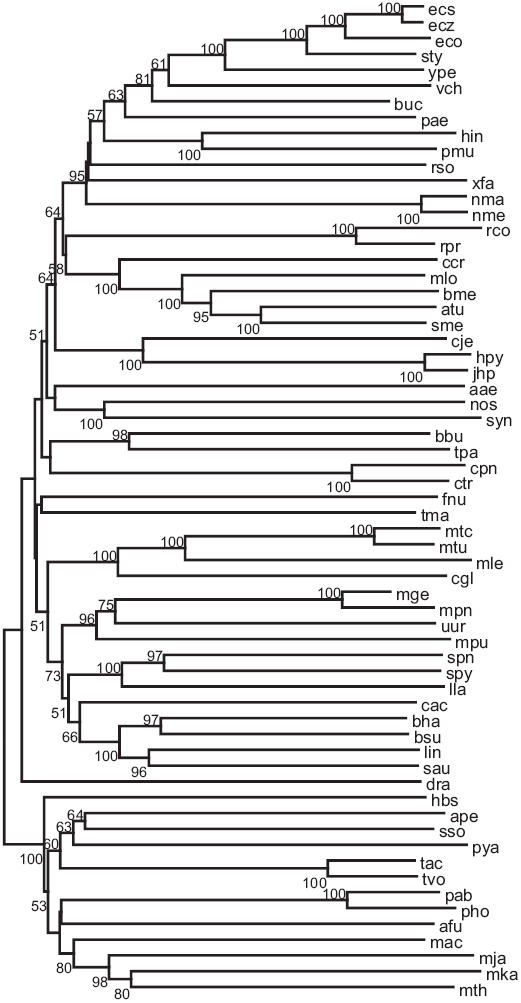
NJ tree inferred from gene bin distance (Tree8). Jackknife support percentages (only if more than 50%) are shown next to each branch. Three-letter species’ abbreviations: **Archaea**: *Archaeoglobus fulgidus* (Afu), *Halobacterium sp.* NRC-1 (Hbs), *Methanosarcina acetivorans* (Mac), *Methanothermobacter* (Mth), *Methanococcus jannaschii* (Mja), *Methanopyrus kandleri* AV19 (Mka), *Thermoplasma acidophilum* (Tac), *Thermoplasma volcanium* (Tvo), *Pyrococcus horikoshii* (Pho), *Pyrococcus abyssi* (Pab), *Pyrobaculum aerophilum* (Pya), *Sulfolobus solfataricus* (Sso), *Aeropyrum pernix* (Ape); **Actinobacteria**: *Corynebacterium glutamicum* (Cgl), *Mycobacterium tuberculosis* H37Rv (Mtu), *Mycobacterium tuberculosis* CDC1551 (MtC), *Mycobacterium leprae* (Mle); γ**-Proteobacteria**: *Escherichia coli* K12 (Eco), *Escherichia coli* O157:H7EDL933 (EcZ), *Escherichia coli* O157:H7 (Ecs), *Yersinia pestis* (Ype), *Salmonella typhimurium* LT2 (Sty*), Buchnera sp.* APS (Buc), *Vibrio cholerae* (Vch), *Pseudomonas aeruginosa* (Pae), *Haemophilus influenzae* (Hin), *Pasteurella multocida* (Pmu), *Xylella fastidiosa* 9a5c (Xfa); α**-Proteobacteria**: *Agrobacterium tumefaciensstrain* C58 (Atu), *Sinorhizobium meliloti* (Sme), *Brucella melitensis* (Bme), *Mesorhizobium loti* (Mlo), *Caulobacter crescentus* CB15 (Ccr), *Rickettsia prowazekii* (Rpr), *Rickettsia conorii* (Rco); **Bacteria**: *Aquifex aeolicus* (Aae), *Thermotoga maritime* (Tma), *Chlamydia trachomatis* (Ctr), *Chlamydophila pneumoniae* (Cpn), *Treponema pallidum* (Tpa), *Borrelia burgdorferi* (Bbu), *Synechocystis* (Syn), *Nostoc sp.*PCC7120 (Nos), *Fusobacterium nucleatum* (Fnu), *Deinococcus radiodurans* (Dra); **Gramplus:** *Clostridium acetobutylicum* (Cac), *Lactococcus lactis* (Lla), *Streptococcus pyogenes* M1GAS (Spy), *Streptococcus pneumoniae* (Spn), *Staphylococcus aureus* N315 (Sau), *Listeria innocua* (Lin), *Bacillus subtilis* (Bsu), *Bacillus halodurans* (Bha), *Ureaplasma urealyticum* (Uur), *Mycoplasma pulmonis* (Mpu), *Mycoplasma pneumoniae* (Mpn), *Mycoplasma genitalium* (Mge); **Proteobacteria:** *Neisseria meningitides* MC58 (Nme), *Neisseria meningitides* Z2491 (NmA), *Ralstonia solanacearum* (Rso), *Helicobacter pylori* 26695 (Hpy), *Helicobacter pylori* J99 (jHp), *Campylobacter jejuni* (Cje).

#### General properties of d_BG_ –based tree

Our tree is in agreement with such well-established notions as the monophyly of each of the two domains, Archaea and Bacteria, and the existence of distinct bacterial clades, such as Cyanobacteria, Proteobacteria, the Termus-Deinococcus group, high-GC and low-GC Gram-positive bacteria. This tree also correctly groups each parasitic species of proteobacteria with its respective free-living relative.

#### Archaea

The topology of this portion of our tree coincides with the Tree5, supporting two clades within archaea, Euryarchaeota and Crenarchaeota-Thermoplasmata. In most genome-based phylogenies, Euryarchaeota are paraphyletic [6, 17]. Only the rRNA tree, Trees3 (Table 1) and tree provided in Fig. 3 in ref. 9, support monophyletic Euryarchaeota that includes Thermoplasmata clade. Thus, our phylogeny argues for monophyly of Euryarchaeota, except for the placement of Thermoplasmata.

*Halobacterium* sometimes is seen as a basal clade in genome-based trees (e.g. Tree6). One plausible explanation of this position is relatively high proportion of genes of bacterial origin in *Halobacterium* [7, 18]. “Bacteria-like” genes in Archaea tend to code for “operational” genes, coding for metabolic enzymes [19, 20], and phylogenies based on sequences of other, “informational” proteins [21] move *Halobacterium* inside euryarchaeota clade. In our phylogeny *Halobacterium* is placed as basal to Euryarchaeota, although with relatively low jackknife support.

#### Proteobacteria

This clade is well-resolved, except that β-proteobacteria *R.solanacearum* and *N.meningitidis* (two strains) intermingle with the γ-proteobacteria. In fact, in rRNA tree, as well as in all genome-based trees with at least three β-proteobacterial species, one or more of them are found within the γ-proteobacteria clade [8, 17]. Several PDS, suggesting HGT between β-proteobacteria and γ-proteobacteria, have been recently pointed out [22]. These PDS include some of the 203 protein families that are conserved in all γ-proteobacteria and resolve to the same topology [23]. γ− and β-proteobacteria frequently share ecological niches, and there is even an example of γ-proteobacteria living symbiotically inside β-proteobacteria, suggesting a lot of opportunity for HGT [22, 24]. Thus, the relationship between γ-proteobacteria and β-proteobacteria appears to be complex and in need of further investigation. α- and ε-proteobacterial clades are both well resolved (high jackknife for every internal branch, Tree8) and placed as basal to the β/γ-proteobacterial cluster.

#### Firmicutes

Our tree provides strong statistical support for the sister status of high-GC and low-GC Gram-positive bacteria (Firmicutes clade). This is the first genome-based tree that supports such hypothesis. Even in Tree5, which is the most similar to our tree in archaeal and proteobacterial clades, high-GC Gram-positive bacteria are joined as one clade with *Deinococcus radiodurans* and cyanobacteria. Monophyly of Gram-positive bacteria has been challenged by analysis of several protein families [25]; nevertheless, it is supported by morphological traits, biochemistry and 16S rRNA tree.

#### Other clades and problem cases

Recently, the existence of novel bacterial clades has been suggested. Several different types of characters and distances support the *Chlamydia*-*Spirochetes* clade [6, 10], and it is seen in our tree, although the statistical support is rather tentative. The radical proposal of the Actinomycetes-Deinococcales-Cyanobacteria clade [6] is not supported in our tree: *Deinococcus radiodurans* appears to be the deepest, but cyanobacteria are placed as basal to all proteobacteria, whereas actinomycetes join other Gram-positive bacteria.

The rRNA tree and several whole-genome studies have resolved *Thermotogales* and *Aquificales* as, respectively, the deepest and second-deepest branch among bacteria [8, 10]. In some trees these bacteria form a clade [6, 16]. In our tree there is no statistical evidence for a specific affinity between the two, *A.aeolicus* being basal to proteobacteria, and *T.maritima* basal to firmicutes. Although both positions have low statistical support, they are not inconsistent with several genome-based trees (Trees4-7, Table 1), including the most recent reconstruction that uses sophisticated methods to take into account PDS [11]. Such placement may also be the most relevant one from biological point of view (see discussion in reference 9).

### Gene Bins and Gene Teams

When our work was in progress, Gene Teams, a rigorous formalization for the concept of “closely placed genes” on two chromosomes, was suggested [26]. Gene Teams approach operates on permutation of genes within a fixed interval over the chromosome. If in both chromosomes the positions of two orthologous genes differ less than given length threshold δ, two genes fall in the same gene δ-set. The maximal δ−set with respect to inclusion constitutes a gene δ-team. This formalism is implemented in a fast TEAM3 software [27], which finds δ-teams using a recursive algorithm.

We used TEAM3 to compute all teams in all possible pairs of 63 microbial genomes within fixed length δ=10. Because TEAM3 does not account for gene duplications (in our case, the COGs that are represented by more than one lineage-specific paralog in the same genome), we retained one such paralogs in each genome and converted the number of genes in teams into the distance measure in the same way as with gene bins. The inferred tree was very similar to our gene bins-based tree, with several minor rearrangements, including branching order in γ-proteobacteria domain, split of the Chlamydia-Spirochetes clade and several others (Figure 3 in additional file 1). Statistical support for most branches was slightly weaker than in the case of gene bins, and, expectedly, global scores were lower since we excluded some genes.

When we restored all gene duplications, however, TEAM3 was no longer able to extract any phylogenetic signal. Now, the highest global pairwise similarity score was found for *Methanosarcina acetivorans* and *Pseudomonas aeruginosa*, two large genomes, which also had the high fraction of duplicate genes (data not shown). We compared distributions of scores obtained by our algorithm and by TEAM3 for genomes with and without lineage-specific duplicates, as well as with genomes in which gene order was reshuffled. As shown in Figure 4 (Supplementary file 2), TEAM3 essentially does not discern between native and shuffled genomes (probability of significant difference between two distributions in Kolmogorov-Smirnov p<0.001). These observations agree with theoretical considerations, suggesting that gene duplication increases the probability of genes occurrence in cluster by chance for window size on the order of 10 (reference [28]; see their Fig. 5 (a) and equation 55 for details).

Recently, the extension of the original Gene Team approach has been proposed, which accounts for the presence of multiple gene copies in a genome [29]. We compared score obtained in this extended approach (*extGT*) and our *Gs* for selected pairs of genomes. *Gs* tends to be similar to respective *extGT* score; for example, *extGT* score of Eco:Hin comparison was 473 (δ=1000), whereas *Gs* score was 467, and comparison Eco:Vch gave *extGT* score 886 (δ=7000) and *Gs* score was 893. Thus, our less formal, empirical gene bin matching procedure is expected to perform comparably to a more rigorously defined extGT approach.

### Horizontal transfer of genes and operons

Phylogenetic discordance between different genes, manifest both at the level of sequence similarity and at the level of presence-absence of orthologs in genomes, is a major source of noises and artifacts in phylogenetic reconstructions. One factor thought to contribute to phylogenetic discordance is horizontal gene transfer [3, 30]. Lawrence [31] furthermore suggested that horizontal transfer of whole operons is more likely to supply the recipient organism with beneficial metabolic functions than transfer of single genes (“selfish operon” hypothesis). The roles of HGT and horizontal operon transfer (HOT) in producing PDS are under ongoing debate [31-33].

We expect our gene-bin distance measure to be resistant to horizontal transfer of single genes: the only way for a singly transferred gene to contribute to *Gs* is if some of its neighbors, within *k*-gene bin, are the same in donor and recipient genomes. On the other hand, HOT, which in principle can transfer whole bins, may cause a more significant increase in *Gs*. The detailed quantitative analysis of both processes remains to be undertaken. In the meantime, we attempted to empirically estimate the contribution of HGT and HOT to our measure of similarity between genomes and to the topology of the phylogenetic tree.

Two cases of likely HGT have received attention: it has been suggested that 246 genes (16% of all genes) in *Aquifex aeolicus* and 450 genes (24%) in *Thermotoga maritima* have been transferred from archaea [34, 35]. We re-evaluated this analysis, using the criteria described in Methods section, and confirmed the HGT hypothesis for 97 genes in *A.aeolicus* and 61 genes in *T.maritima* genomes, respectively. We then used a recent compilation of putative HGT events throughout 41 genomes [32], and found that only 4 of 97 genes in *A.aeolicus*, and 17 of 67 genes in *T.maritima.* When all these putative horizontally transferred genes were removed from our dataset, only a slight decrease in jackknife support was observed, but there were no topological changes in the phylogenetic tree. It is also notable that the *A.aeolicus* and *T.maritima* clades appear to be more recent evolutionary events in our tree than in Trees1-3 and 6, further arguing that the HGT/HOT has little effect on their position relative to archaea ^1^.

### Discussion

Gene order in bacterial genomes is eroded by recombination and gene gain/loss. Over short evolutionary distances, such as those between two species of *Chlamydia*, or two strains of *H. pylori*, chromosomal order of genes is essentially the same. Genome rearrangements in closely related species often preserve local gene order too, by way of translocation of large DNA fragments, often symmetrically with regards to the replication origin [37, 38]. Further apart in evolution, the picture is different: for example, within one subdivision of proteobacteria, *H. influenzae* shares most genes with *E.coli*, but only short gene strings, albeit many of them, are conserved in the two genomes. Presumably, this is because extensive gene loss in *H. influenzae* resulted in jumbling of its genome [39]. Finally, at extremely large evolutionary distances, such as between bacteria and archaea, there has been ample time for all types of genome shuffling, so that a few strings of genes, typically coding for the stoichiometric components of the same molecular complex, are conserved [12, 40].

These observations suggest that (dis)similarity of gene order in two species can reflect the time since their divergence [13, 41, 42] and may therefore be useful for reconstruction of evolutionary events. Automated approaches have been proposed for finding perfect strings of gene colinearity and to account for occasional indels [13, 43], and criteria of statistical significance for perfectly matched strings have been recently generalized for the case of approximate matches [28]. Thus far, however, analysis of gene order has been done primarily with the aim of finding functionally associated genes by their proximity in the genome, whereas the efforts of phylogeny reconstruction on the basis of gene order focused mostly on conservation of adjoining gene pairs [6, 8].

A special case of phylogeny reconstruction from gene order is the study of ordered gene permutations and reversal distances, i.e., the minimum number of chromosomal inversions required to convert one gene order into another (reviewed in [44] and [45]). These methods are highly relevant to the analysis of organelle genomes and those of certain DNA viruses, which share stable sets of orthologs and evolve mostly by such inversions. In contrast, microbial genome evolution is not dominated by permutations of a constant gene set; instead, gene gains and losses play a major role, so that there are only about 80 universally conserved genes in prokaryotes [46]. Here, we propose a measure of evolutionary distance based on local gene conservation in a broad sense, in which indels and local permutations of gene order are tolerated.

Very recently, the methodology underlying the computation of edit distance between two genomes was extended to take into account not only inversions of chromosome segments, but also gene duplications and deletions [47]. This type of rigorously defined distance measure helps to infer correct phylogeny between closely related gamma proteobacteria [48], however, its performance with distantly related genomes has not been assessed.

Gene bin distance proposed here correctly resolves several clades that contain both free-living and parasitic bacteria, with almost an order-of-magnitude difference in the number of genes, indicating that our measure is not very sensitive to gene loss. Evidently, strong phylogenetic signal remains captured by the local gene content, even as other genes are deleted from the genome. There is also an indication that HGT and HOT have relatively small impact on our distance measure, possibly arguing that horizontal transfer, when it occurs, is not strongly associated with local gene order conservation, at least in the specific cases examined her (i.e., inter-domain transfers between archaea and bacteria). Remarkably, the informal measure of local gene order conservation that we adopted in this work allowed us to discern substantial phylogenetic signal, on a par with the performance of several more rigorously defined measures of local gene order conservation. We did not attempt to optimize the *k* parameter in this study, and may be missing some of the evolutionary signal because of that. Moreover, the COG database, which we used as a source of gene content information, includes only genes present in three or more clades. Addition of genes shared by two genomes, may allow one to produce even more robust phylogenies on the basis of local conservation of gene order.

## Methods

### Data set

Genome content of 66 microbial species is summarized in the COG database at NCBI (http://www.ncbi.nlm.nih.gov/COG/new). There were 4873 COGs from 66 complete genomes of unicellular organisms in the COG database, as of early 2004 [49]. After excluding 285 fungi-specific COGs, we have 4588 COGs from 63 prokaryotes. The information about the linear order of COGs in bacterial and archaeal chromosomes was retrieved from the Genomes division of GenBank. COGs locations in microbial genomes were converted into 63 gene order vectors, where each coordinate (from 484 dimensions for *Mycoplasma genitalium* to 6746 for *Mesorhizobium loti*) is represented either by the appropriate COG identifier, or by a blank, if a gene does not belong to any COG in the database. A COG can appear more than once in the same genome, because the algorithm of COGs database construction sometimes treats lineage-specific gene duplications as one COG [50].

### Comparison of gene bins in two genomes

We are looking for the optimal assignment of bins (pair of chromosomal segments from two genomes, containing from 0 to k orthologous genes). The optimality means that we want to assign each bin in a smaller genome to a bin in a larger genome, in order to maximize the global assignment score *Gs*, which is the sum of local assignment scores *s*_*ij*_, i.e., the number of orthologous genes in two bins. One issue that complicates the optimization is that going after the highest *s*_*ij*_ does not guarantee maximal *Gs*. Consider two genomes, each with two non-overlapping bins of length *k*=10 (this value was used throughout the study as a compromise between the ability to compare groups of genes at or above the operon level and the decline of global scores as *k* becomes larger) and the following set of *s*_*ij*_: *s*_*11*_=6; *s*_*12*_ =4; *s*_*21*_ =5; *s*_*22*_ =0. Any algorithm that starts with examination of the bin(1,1) and seeks the highest *s*_*ij*_, will first assign bin(1,1) to bin (2,1), and then assign bin(1,2) to bin(2,2). The global score *Gs*=*s*_*11*_+*s*_*22*_ will be 6, while in the alternative assignment, *Gs*=*s*_*12*_+*s*_*21*_ would be 9. The other problem is how to break ties.

We designed a fast heuristic algorithm, which finds a global assignment in an iterative mode. The complete set of possible assignments contains at each iteration *nm* elements, where *n* and *m* are, respectively, the numbers of unassigned bins in smaller and larger genomes. One idea is to reduce this list of candidate assignments to 2*n*, by finding, for each segment *i* in the smaller genome, two bins in the larger genome, those with the highest and second-highest *s*_*ij*_. We assume that these two scores capture most of local gene order conservation, i.e., that the contents of a bin can be split between two bins of the same length in another genome, but further splits cause rapid decay in *s*_*ij*_ and do not contribute much to Gs. The bin pair is selected among the still unassigned pairs by maximizing the difference between the two highest bin scores: max_*i*_{*s*_*ib*_1__ − 0.5*s*_*ib*_2__}, where *s*_*ib*_1__ = max_*j*_{*s*_*ij*_} and *s*_*ib*_2__ = max_*j*_{*s*_*ij*_} ≤ *s*_*ib*_1__. In the aforementioned example, the optimal assignment matches bin(1,1) to bin(2,2) and bin(1,2) to bin(2,1). The algorithm described below produces a total score of 9 as follows: for the first bin *s*_1*b*_1__ and *s*_1*b*_2__ are 6 and 4, for the second bin *s*_2*b*_1__ and *s*_2*b*_2__ are 5 and 0 and 6-0.5*4=4 is less than 5-0*2=5. The ties are broken deterministically. Our second time-saving measure is to assign one bin at a time, starting with the highest-scoring bins. When a bin is assigned, it is removed from further examination, reducing the number of remaining bins.

Although we assumed the non-overlapping bins, it is in fact more natural to consider the overlapping genome segments (sliding windows), to avoid arbitrary split in the middle of a high-scoring bin. With sliding windows, the initial number of bins, *n*=*N/k* (where *N* is the number of genes in the genome), becomes *N-k+1*, but the algorithm is almost the same, with just one extra step, namely that after each bin assignment, all bins that overlap the assigned bin are removed from further consideration.

The formal description of the algorithm is as follows:

1. Divide pair of genomes *G*_*i*_, *G*_*j*_ containing *N*_*i*_, *N*_*j*_ genes into, respectively, *N*_*i*_-*k*+1 and *N*_*j*_-*k*+1 *k*-gene bins.
2. Compute rectangular matrix (*N*_*i*_-*k*+1)x(*N*_*j*_-*k*+1) **S** of bin scores **S**={*s*_*ij*_}. *s*_*ij*_ is the number of genes (COGs) that have members in two bins, one in each genome: max *s*_*ij*_ = *k*. Only one appearance of gene in a bin is counted, i.e., local gene duplications are ignored.
3. Find optimal assignment of bins in a pair of genomes *G*_*i*_, *G*_*j*_, steps 3.1-3.6.
  3.1 If there are unassigned bins in smaller genome, do 3.2-3.4.
  3.2 For all unassigned bins in smaller genome, find first highest scoring match (HSM) and write down the index of the corresponding bin in larger genome.
  3.3 For all unassigned bins in smaller genome, find second HSM.
  3.4 Find bin in smaller genome, which maximizes (HSM_1_-0.5*HSM_2_)
  3.5 Assign this bin *i* in the smaller genome to bin *j* in the larger genome. Mark these bin as assigned.
  3.6 Mark as assigned all the bins that overlap bins *i, j* and go back to 3.1.
4. Compute the global assignment score *Gs*_*ij*_ as the sum of bin scores, *s*_*ij*_ of assigned bins.

### Test for gene order preservation among horizontally transferred genes

Bacterium *Aquifex aeolicus* has many genes that appear to be more similar to archaeal orthologs than to bacterial ones [34, 51]. We use phylogenetic criteria to test the HGT hypothesis for each gene in *A.aeolicus* and accepting it when one of the two was true:

- gene was found only in *A.aeolicus* and archaea, or in *T.maritima*, *A.aeolicus* and archaea.
- gene was found in other bacteria besides *A.aeolicus*, and the *A.aeolicus* gene was grouped with archaeal genes in a phylogenetic tree.

To test the second proposition, we aligned each of the *A.aeolicus* genes to its homologs from other species using DIALIGN [52] or CLUSTALX [53] programs, and reconstructed NJ trees using Poisson-corrected gamma distance in MEGA program [54]. If there was more than one *A.aeolicus* gene in a COG, we accepted HGT hypothesis when all *A.aeolicus* genes agreed with one of these conditions. Similar criteria were applied to detect HGT in *T.maritima* [35], substituting *T.maritima* for *A.aeolicus* in rule 2.

## Additional data files

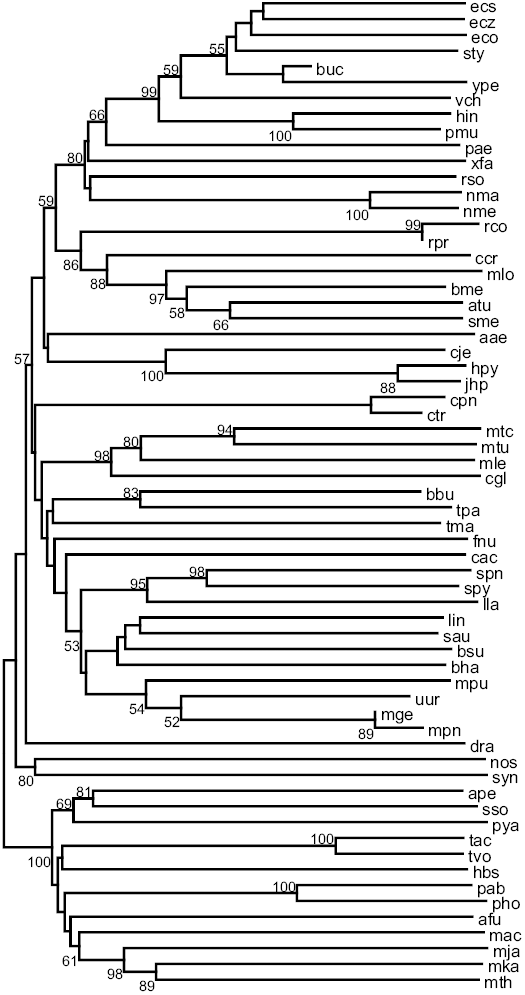
Additional data file 1: SupFig3.pdf. Supplementary Figure 3. NJ tree inferred from gene team distance. Jackknife support percentages were computed the same way as for tree in Fig.2.

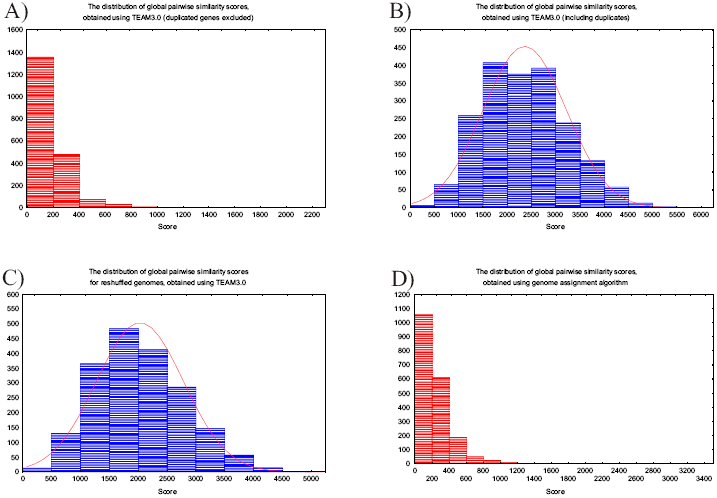
Additional data file 2: SupFig4.pdf. Supplementary Figure 4. The distributions of global pairwise similarity scores.

### Additional files provided with this submission

**Additional file 1: SupFig3.pdf: 24KB**

http://www.biomedcentral.com/imedia/1245219327532973/sup1.pdf

**Additional file 2: SupFig4.pdf: 53KB**

http://www.biomedcentral.com/imedia/2090400688488743/sup2.pdf

## Footnotes

1 Note that the alternative evolutionary explanation for “archaea-like” genes in *Aquifex* has been put forward, such as their origin in the common ancestor of Bacteria and Archaea followed by massive gene loss in nearly all bacterial lineages [36]. Under this scenario, no HGT has happened. Whatever the true history of these genes is, it is inconsequential for our reconstruction: indeed, our criteria err on the side of adding the HGT events, yet our conclusion is that HGT does not affect the trees built with our distance measure.

